# Functional networks in the infant brain during sleep and wake states

**DOI:** 10.1101/2023.02.15.528718

**Authors:** Tristan S. Yates, Cameron T. Ellis, Nicholas B. Turk-Browne

## Abstract

Functional brain networks are assessed differently early in development than at maturity: infants are almost universally scanned during sleep, whereas adults are typically scanned awake while resting or performing tasks. Observed differences between infant and adult functional networks may thus reflect these differing states of consciousness rather than or in addition to developmental changes. We explore this question by comparing functional networks in fMRI scans acquired from infants during natural sleep and awake movie-watching. As a reference, we also acquired fMRI scans in adults during awake rest and awake movie-watching. Whole-brain functional connectivity was more similar within-state (sleep-sleep, wake-wake) than across-state (sleep-wake) in infants, demonstrating that movies elicit a different network configuration than typical sleep acquisitions. Indeed, a classifier trained on patterns of functional connectivity during infant sleep versus wake robustly decoded the state of additional infants and even generalized to decode rest versus movie in adults; interestingly, a classifier trained on rest versus movie in adults did not generalize nearly as well to sleep versus wake in infants. Moreover, the overall level of similarity between infant and adult functional connectivity was modulated by adult state (stronger for movie than rest) but not infant state (equivalent for sleep and wake). Nevertheless, the network connections that drove similarity between infants and adults, particularly in frontoparietal network, were modulated by infant state. In sum, infant functional connectivity can differ between sleep and wake states, highlighting the potential value of awake data for studying the early development of functional brain networks.

**Significance statement:** Functional networks in the infant brain provide a foundation for early cognitive abilities and act as a marker of brain maturation and developmental disorders. What we know about these networks comes from fMRI data acquired during sleep, given the challenges of awake infant fMRI. This contrasts with the dominant approach in older populations of assessing networks during awake rest or tasks. These differing levels of consciousness cloud the interpretation of developmental changes. Here we show that whole-brain functional connectivity differs between sleeping and awake infants, and that the similarity of these infant states to adults loads on dissociable network connections. This research suggests that a full understanding of early functional brain networks will benefit from complementary insights in awake infants.

## Introduction

The discovery of resting-state functional connectivity, or the synchronous fluctuation of brain regions during rest, transformed the way that neuroscientists think about brain function (Biswal et al., 1995; Biswal, 2012). In the decades since its discovery, resting-state fMRI has been used to map different functional networks of the brain (e.g., default mode, control, and salience networks; Raichle et al. 2001; Dosenbach et al. 2006; Corbetta and Shulman 2002), describe brain network properties like modularity and flexibility (Bullmore and Sporns, 2009; Bassett and Sporns, 2017), predict cognitive abilities (Finn et al., 2015; Rosenberg et al., 2016), characterize reliable individual differences (Gordon et al., 2017; Gratton et al., 2020), and establish clinical biomarkers (Baker et al., 2019; Dhamala et al., 2022). Given this, resting-state fMRI has been increasingly used in developmental populations to track the emergence of functional networks relevant to cognition (Grayson and Fair, 2017; Fair et al., 2021) and developmental disorders (Milham et al., 2012; Finn et al., 2014; Hull et al., 2017), including dozens of published studies in infants (Gao et al., 2017; Zhang et al., 2019) and large-scale, longitudinal data collection efforts (e.g., ABCD and HBN in adolescents, dHCP, BCP, and HBCD in infants; Casey et al. 2018; Alexander et al. 2017; Fitzgibbon et al. 2020; Eyre et al. 2021; Howell et al. 2019; Volkow et al. 2021).

Resting-state functional connectivity is typically acquired while participants perform the simple task of staring at a fixation cross and letting their minds freely wander. Participants are actively discouraged from closing their eyes and falling asleep, as this impacts functional network measurements (Tagliazucchi and Laufs, 2014). More recently, however, naturalistic stimuli (such as movies) have become popular for collecting functional network measures (Sonkusare et al., 2019), particularly in developmental populations (Vanderwal et al., 2019), given that they help reduce head motion (Vanderwal et al., 2015; Frew et al., 2022). Functional brain networks differ between restand movies in both adults (Betti et al., 2013; Lynch et al., 2018) and over development (Sanchez-Alonso et al., 2021). In fact, movies are increasingly being recognized as better than traditional fixation-rest tasks for test-retest reliability (Wang et al., 2017; Zhang et al., 2022), characterizing individuals (Vanderwal et al., 2017), and predicting behavior (Finn and Bandettini, 2021; Gal et al., 2022). This has led some to argue for the need to examine the dynamics of cognitive states during typical resting-state (Gonzalez-Castillo et al., 2021) and to embrace more task-like paradigms (Finn, 2021).

Yet, when it comes to infants, resting-state functional connectivity is measured quite differently: infant functional networks are almost exclusively assessed during natural sleep (Zhang et al., 2019). This is a reasonable approach, especially for young infants who spend lots of time in sleep states (Poppe et al., 2021). Moreover, from a practical perspective, it is notoriously difficult to collect fMRI data in awake infants, who move at will, cannot understand or follow instructions, have short attention spans, and need frequent touch, feeding, and diaper changes. Nonetheless, studying the infant brain only in the sleep state may provide an incomplete picture, given sleep/wake differences in adults (Tagliazucchi and Laufs, 2014; Song and Tagliazucchi, 2020) and the sometimes limited reliability of functional connectivity in sleeping infants (Dufford et al., 2021; Wang et al., 2021). Thus, apparent differences in functional brain networks between infants and older populations may not be attributable entirely to development *per se* and could be confounded by different states of consciousness. In line with this, the organization and properties of infant functional networks during sleep are more similar to adults in deep sleep than adults who are awake (Mitra et al., 2017).

It is unknown whether scanning infants in an awake state may yield meaningful differences in functional network measures. The small number of published awake infant fMRI studies contain a suggestive example of how sleep/wake state can matter: the prefrontal cortex of infants is activated by forward versus backward speech during wake but not sleep states (Dehaene-Lambertz, 2002). Indeed, despite undergoing protracted structural development, the prefrontal cortex contributes to cognitive function early in life (Raz and Saxe, 2020; Ellis et al., 2021). From this example, higher-order associative networks, including nodes in frontal areas, may be more engaged and appear more functionally mature when infants are awake compared to asleep.

Recent advances have made it possible to scan infants with fMRI while they are awake and engaged in cognitive tasks (Ellis et al., 2020). This allowed us to examine infant functional networks during the wake state. Namely, we measured functional connectivity in infants scanned while they watched a movie awake and compared this to functional connectivity while they slept naturally. We further compared the infant data to awake adults scanned while watching the same movies or completing a canonical resting task with fixation. After parcellating the brain (Schaefer et al., 2018), we first tested whether infants have a more similar pattern of whole-brain functional connectivity within (wake-wake, sleep-sleep) versus across (wake-sleep) behavioral states. We next used pattern classification to decode behavioral state from functional connectivity patterns within and across age groups. We then tested whether having infants watch movies increases the similarity of their functional connectivity patterns to adults (who are typically scanned awake in functional connectivity studies). To interpret the resulting similarity, we quantified the contributions of individual network connections within and across functional networks. The results show that although both infant sleep and wake yield adult-like functional connectivity, the networks involved and how they are configured is modulated by state. This highlights the value of both sleep and awake infant fMRI for characterizing the nature and early development of functional brain networks.

## Methods

### Participants

Sleep fMRI data were collected from 14 unique infants (7 female) who fell asleep naturally while we collected data for other awake fMRI experiments (not discussed here). In total, we acquired 20 sleeping runs from infants ranging from 3.9 to 24.9 months of age (*M* =11.2, *SD* = 5.0 months). One participant contributed two sleep runs in the same session, separated by 8.5 minutes. Five participants had more than one session with usable sleep data (*M* = 1.4 sessions, range: 1 to 3 sessions), with an average of 5.3 months between sessions (range: 1.3 to 15.0 months). One additional sleep run was excluded because the infant woke up after 1.5 minutes.

Movie-watching fMRI data were collected from 22 unique infants (14 female) who watched one of two cartoon movies, described in detail in a previous publication (Yates et al., 2022). In total, we acquired 34 movie-watching runs from infants ranging from 3.6 to 32.6 months of age (*M* = 12.8, *SD* = 7.3 months). Three participants had two usable movie-watching runs in the same session, separated by an average of 10.0 minutes (range: 3.8 to 15.9 minutes). Six infants completed more than one session with usable movie-watching runs (*M* =1.5 sessions, range: 1 to 6 sessions), with an average of 3.5 months between consecutive sessions (range: 1.4 to 6.3 months). The 34 runs do not include data from infants who had excessive head motion (>3mm framewise displacement) for more than 4% of the time (N = 37), who did not look at the screen during more than half of the movie (N = 6), who did not complete the movie because of fussiness (N = 9), or because of technical error (N = 1). The strict threshold of 4% maximum motion TRs (rather than 50%, which we typically use for task-based awake infant fMRI studies) was chosen to equate the average proportion of usable TRs between infant sleep and infant wake groups.

For comparison, we collected awake resting (staring at fixation) and movie-watching runs from 12 adults (7 female) aged 18 to 32 years (*M* = 21.42, *SD* = 3.40 years). We supplemented these 12 new adult moviewatching runs with 66 additional movie-watching runs we previously collected for other studies from 48 unique adults (27 female, age: *M* = 21.94, *SD* = 3.18 years). In total, we thus had 12 runs of awake fixation-rest and 78 runs of movie-watching from adults. This does not include movie runs in which adults fell asleep part-way (N = 2) or runs with excessive head motion (N = 1).

The study was approved by the Human Subjects Committee (HSC) at Yale University. All adults provided informed consent, and parents provided informed consent on behalf of their infant.

### Materials

Infant sleep runs were collected during natural sleep with the visual display turned off or dimmed. Sleep state was not assessed physiologically, but all infants were assumed to be asleep based on extended eye closure and stillness, as viewed online via an MR-safe camera. Infants stayed asleep for variable durations, between 1.76 and 5.97 minutes (*M* = 4.31, *SD* = 1.34 minutes). We often stopped a sleep scan after 5 minutes to transition to anatomical scans or to try waking the child up for additional awake functional runs. Otherwise, we kept collecting sleep data until the child naturally woke up.

Infant movie runs were collected while infants watched one of two silent cartoon movies, previously described in Yates et al. (2022). The first movie, “Aeronaut”, is a 3-minute long segment of a short film about a miniature pilot and a little girl (https://vimeo.com/148198462). For all participants, the movie spanned 45.5 visual degrees in width and 22.5 visual degrees in height. The second movie, “Mickey”, is a 2.37-minute long segment of popular cartoon show where characters celebrate a birthday party. This movie was displayed in a smaller size, spanning 22.75 visual degrees in width and 12.75 visual degrees in height. Movie runs were collected while infants watched Aeronaut once (N = 25), Aeronaut twice in a row (N = 1), Mickey once (N = 7), or Mickey twice in a row (N = 1). Infant movie runs were therefore between 2.37 and 6.00 minutes long (*M* = 3.01, *SD* = 0.66 minutes).

Adult fixation-rest runs were collected during quiet rest while participants stared at a white fixation cross (2 visual degrees) on a black background. Participants were not instructed to think about anything in particular. These rest runs always lasted 5 minutes, and the order of rest and movie-watching runs was counterbalanced across the 12 participants who completed both. Adult movie-watching data consisted of participants who watched Aeronaut once (N = 52), Mickey once (N = 11), or Mickey twice in a row (N = 15), often interleaved with other movie-watching or experimental runs not described here. Adult movie runs were between 2.37 and 4.73 minutes long (*M* = 3.24; *SD* = 0.76 minutes).

The code used to display the movies and fixation is available at https://github.com/ntblab/experiment_menu/tree/Movies/. The code used to perform the data analyses is available at https://github.com/ntblab/infant_neuropipe/tree/RestingState/. Raw and preprocessed functional data and anatomical images will be released publicly upon publication.

### Data acquisition

We used a previously validated procedure for collecting infant fMRI data (Ellis et al., 2020) and adult comparison data. All adult data and most infant data (N = 16 infant sleep runs, N = 26 infant movie runs) were collected at the Brain Imaging Center in the Faculty of Arts and Sciences at Yale University. Data were acquired using a Siemens Prisma (3T) MRI using the bottom half of the 20-channel head coil. We used a whole-brain T2* gradient-echo EPI sequence (TR = 2s, TE = 30ms, flip angle = 71, matrix = 64×64, slices = 34, resolution = 3mm iso, interleaved slice acquisition) to acquire functional images for both adults and infants. For infants, we collected a T1 PETRA sequence (TR1 = 3.32ms, TR2 = 2250ms, TE = 0.07ms, flip angle = 6, matrix = 320×320, slices = 320, resolution = 0.94mm iso, radial slices = 30000) as the anatomical image. For adults, we collected a T1 MPRAGE sequence (TR = 2300ms, TE = 2.96ms, TI = 900ms, flip angle = 9, iPAT = 2, slices = 176, matrix = 256×256, resolution = 1.0mm iso), which included the top half of the 20-channel head coil. The remaining infant movie and sleep runs were collected at the Scully Center at Princeton University (N = 4 sleep, N = 5 movie) using a Siemens Skyra (3T) MRI and at the Magnetic Resonance Research Center (MRRC) at Yale University (N = 3 movie) using a Siemans Prisma (3T) MRI. All acquisition procedures were the same, with the exception that the functional EPI sequence had slightly different parameters at these latter two sites (TE = 28ms, slices = 36).

### Procedure

All procedures followed lab conventions that have been used in our prior publications (e.g., Ellis et al., 2021; Yates et al., 2022). Infants and their parents met with researchers prior to their first scanning session. For almost all participants, these visits were conducted in-person as a “mock scanning” session; however, some visits were conducted over zoom in accordance with COVID-19 policies. We then scheduled scans for times the families thought their infant would be most comfortable. We extensively screened parents and infants for metal before and on the day of the scan. Infants were then equipped with three layers of hearing protection: silicone inner ear putty, over-ear adhesive covers, and ear muffs. Parents were permitted to bring comfort items (e.g., metal-free blankets) for their infant, and infants were wrapped with a vacuum pillow to reduce movement. We projected stimuli directly on the ceiling surface of the scanner bore, and recorded participants’faces with a camera (MRC high-resolution camera) during the session. Adults similarly viewed stimuli on the scanner bore ceiling and were monitored with a camera, but only had two layers of hearing protection (earplugs and optoacoustics noise-canceling headphones), were not given comfort items or a vacuum pillow, and did not attend a mock scanning session. For both adults and infants, additional tasks were sometimes run during their scanning session.

### Gaze coding

During infant movie-watching runs, gaze was coded offline by 2-6 coders (*M* = 2.39, *SD* = 0.98). Coders determined whether the participant’s eyes were on-screen, off-screen (i.e., closed, blinking, or looking off of the screen), or undetected (i.e., out of the camera’s field of view). In one infant, technical issues prohibited us from collecting gaze data, but this infant was monitored live by a researcher during data collection and determined to be attentive enough to warrant inclusion. Coders were highly reliable, reporting the same response code on an average of 93.76% (*SD* = 4.84%; range across participants = 76.62–99.62%) of frames. The modal response across coders from a moving window of five frames was used to determine the final response for the frame centered in that window. The response from the previous frame was used in the case of ties. Frames were pooled within TRs, and the average proportion of TRs included for eyes being on-screen was high (*M* = 92.75%, *SD* = 9.46%; range across participants = 63.38–100%). Gaze data were not collected during infant sleep or adult rest runs, and gaze data were not analyzed for adult movie runs.

### Preprocessing

Data were preprocessed using a custom awake infant fMRI pipeline (Ellis et al., 2020). All adult data came from distinct functional runs, while movie and sleep data from infants were sometimes cleaved into pseudoruns when another task was performed in the same functional run (N = 14 sleep runs, N = 20 movie runs). We discarded three burn-in volumes from the beginning of each run/pseudo-run. Then we determined the centroid volume of each run/pseudorun by calculating the Euclidean distance between the brain mass in all volumes and choosing the volume that minimized the spatial distance to all other volumes, using this as the reference for motion correction. Volumes were realigned using slice-timing correction. During preprocessing, we excluded timepoints with greater than 3 mm of translational motion, temporally interpolating them so as not to bias linear detrending. In the current study, we only included participants with more than 96% of TRs that were not motion TRs; thus, almost all timepoints were included after motion exclusion in: infant sleep runs (*M* = 99.74%, *SD* = 0.75%; range across participants = 96.72–100%), infant movie runs (*M* = 99.64%, *SD* = 0.76%; range across participants = 96.67–100%), adult rest runs (all 100% usable), and adult movie runs (*M* = 99.99%, *SD* = 0.12%; range across participants = 98.89-100%). In subsequent analyses, we excluded these motion TRs, and for infant movie runs, we also excluded timepoints during which eyes were closed for a majority of movie frames in the volume (out of 48, given the 2-s TR and movie frame rate of 24 frames-per-second). We constructed the mask of brain versus non-brain voxels by thresholding based on the signal-to-fluctuating-noise ratio (SFNR) (Friedman and Glover, 2006). Then, data were spatially smoothed with a Gaussian kernel (5mm FWHM) and linearly detrended in time. We used AFNI’s (https://afni.nimh.nih.gov) despiking algorithm to attenuate aberrant timepoints within voxels. Finally, after removing excess burn-out TRs, functional data were *z*-scored within run/pseudorun.

The centroid functional volume was first registered to the anatomical image using FLIRT in FSL (Jenkinson et al., 2012), and adjusted manually as needed using MR-Align from mrTools (Gardner lab). Then, the anatomical image was aligned into standard space using ANTs (Avants et al., 2011), a non-linear alignment algorithm. For infants, we used an initial linear alignment with 12 DOF to align their anatomical data to an age-specific infant template in MNI space (Fonov et al., 2009), followed by non-linear warping using diffeomorphic symmetric normalization. After this alignment, we used a predefined transformation (12 DOF) to linearly align between the infant template and adult MNI standard (MNI152). For adults, we used the same alignment procedure, except participants were directly aligned to the adult MNI standard. For all analyses, we only considered voxels included in the intersection of all infant and adult brain masks.

### Whole-brain functional connectivity

Functional connectivity matrices were created using the Schaefer brain atlas parcellation (Schaefer et al., 2018). The Schaefer atlas consists of parcels discovered from resting-state functional connectivity data in adults and is available at multiple resolutions. To match the number of brain regions found in neonatal resting-state analyses (Scheinost et al., 2016), and to account for potential anatomical variability across participants, we used the 100-parcel version of the Schaefer atlas. These parcels were matched to 7 functional networks — visual, somatomotor, dorsal attention, ventral attention, limbic, frontoparietal control, and default (Yeo et al., 2011). The number of parcels that made up each network ranged from 5 (limbic) to 24 (default), with an average of 14.29 parcels. Note that these network labels for parcels correspond to *adult* functional networks, and their applicability to infant functional networks has not been established. Nonetheless, we use these labels throughout the manuscript to give an idea of the localization of our effects with respect to adult data.

To construct individual functional connectivity matrices, we averaged BOLD activity over all voxels in each parcel and correlated this average timeseries with every other parcel using Pearson correlation, after excising high-motion timepoints. The resulting coefficients were transformed into *z*-scores by normalizing to the average and standard deviation of the correlations across parcels, to account for potential differences in absolute correlation values across groups. We created group-level connectivity matrices by averaging across runs within group. To measure similarity within group, we correlated the upper triangle of each individual’s functional connectivity matrix with the average functional connectivity matrix for all but that run. We used an analogous procedure for calculating similarity across groups, by correlating an individual’s upper triangle with the other group’s average of all runs.

We used bootstrap resampling to evaluate the statistical reliability of these functional connectivity similarity scores (Efron and Tibshirani, 1986). Specifically, we randomly sampled with replacement from the run-level similarity values (*z*-scored Pearson correlations) to form a new sample of the same size as the original group, computed the average similarity of the sampled values, and repeated this procedure 1,000 times to create a sampling distribution. This approach also allowed us to compute the reliability of differences between states and groups: on each resampling iteration, we subtracted the mean similarity within one state or group from the mean similarity with another state or group to create a sampling distribution of the difference. We calculated *p*-values as the proportion of iterations with the opposite sign of the original effect, doubled for a two-tailed test.

### Decoding state within and across age groups

We further investigated how states modulated functional connectivity using pattern classification. The input features for the classifier models were the correlation values for every parcel-to-parcel connection in the upper triangle of the functional connectivity matrix and the output was the state during which functional connectivity was measured. To assess how different networks contributed to decoding accuracy, we also learned and evaluated classifiers on the subset of features within and across specific networks.

For each age group, we divided runs into training and test sets (approximately 90% training and 10% test), while subsampling runs from the more populous state so that the classifier was trained and tested with 50% of the examples from each state. In an additional analysis, we retained the original distribution of participants from each state (e.g., 63% of infant runs were awake movie-watching, so the training and test set each had 63% movie runs; Figure S3). In the training set, we further split the data to tune the cost parameter of a linear support vector machine classifier. The best cost parameter from these inner loops was used to train the classifier on the whole training set that was then applied to the held-out test data. This procedure was iterated across 10 folds. We used a generalization approach to assess whether the same features were important for classifying state in adults and infants by applying the best classifier trained in one age group to the other age group.

To determine statistical significance for classification analyses, we used bootstrap resampling. The *p*-value was calculated as the proportion of iterations where classification accuracy was lower than chance (50%), doubled for a two-tailed test. For the control analysis, we compared the true accuracy to a permuted null distribution. Specifically, we repeated the full classification pipeline 1,000 times after randomly shuffling the labels. We then transformed the true classification accuracy into a *z*-score by subtracting the mean of the null distribution and dividing by its standard deviation. The *p*-value was calculated as the proportion of iterations in the null distribution with higher classification accuracy than the true effect, doubled for a two-tailed test. For network analyses, we visualize all network connections that are significant at the 0.05 level, and additionally indicate which connections survive Holms-Bonferroni correction on the alpha value.

### Similarity of functional connectivity between infants and adults

We next tested for differences in the similarity of functional connectivity across groups (e.g., infant sleep to adult movie vs. infant movie to adult movie). First, we tested for the main effect of infant state on overall similarity to adults: (infant sleep to adult movie + infant sleep to adult rest) – (infant movie to adult movie + infant movie to adult rest). Then, we tested for the main effect of adult state on overall similarity to infants: (infant sleep to adult rest + infant movie to adult rest) – (infant sleep to adult movie + infant movie to adult movie). Finally, we tested for an infant state by adult state interaction: (infant sleep to adult movie – infant movie to adult movie) - (infant sleep to adult rest – infant movie to adult rest). For each test, we performed bootstrap resampling on the contrast values across participants to determine significance.

Finally, we evaluated the relationship between functional connectivity similarity and participant age with bootstrap resampling. Namely, we randomly sampled bivariate similarity-age pairs with replacement and re-calculated the Pearson correlation between similarity and age over the sampled pairs on each of 1,000 iterations. Again, the *p*-value was calculated as the proportion of resampled coefficients with the opposite sign as the original effect, doubled for a two-tailed test.

### Contribution of individual connections to overall network similarity

To understand which parcels drove overall similarity between two functional connectivity matrices, we made use of the fact that the Pearson correlation between two variables (in this case, vectorized matrices) is the sum of their pointwise products, after normalizing each variable by mean-centering and dividing by the root sum of squares (Turk-Browne, 2013). Thus, for each run, the normalized pointwise product for a given cell in the matrix quantifies how much the functional connectivity between that pair of parcels contributed to the overall network similarity between that run and the average of other runs. We tested where in the brain these normalized pointwise product values differed between group and state comparisons. For example, we asked whether the same or different connections made sleeping vs. awake infants similar to adults.

First, for each comparison of functional connectivity between an individual run and the average of a different group (e.g., a single infant sleep run to all adult rest runs), we created a normalized pointwise product matrix, where summing the values of the upper triangle of this matrix would equal the Pearson correlation between the upper triangles of the individual participant and group functional connectivity matrices. All values of pointwise product matrix were then converted to relative or proportional scores by dividing them by the overall correlation. We visualize the magnitude of these parcel-by-parcel contributions to the overall similarity between two groups by averaging across participants and plotting the top 1% of contributing connections on a Circos plot.

Next, we tested for differences in the contributions of connections between one group comparison (e.g., infant sleep and adult rest) and another group comparison (e.g., infant movie and adult rest). To simplify, we averaged normalized pointwise product values for each group comparison at the level of networks (e.g., all parcel connections within the visual network, all parcel connections between the visual and somatomotor networks). Then, as before, we used bootstrap resampling to calculate the statistical reliability of differences. On each iteration, we subtracted the mean normalized pointwise product of one group comparison from the mean normalized pointwise product value of the other group comparison. In additional analyses, we also used a connection *lesioning* approach, where we assessed how removing one connection at a time changed the resulting correlation between groups (Figure S4). As with the decoding analysis, we visualize all network connections that are significant at the 0.05 level, and additionally indicate which connections survive Holms-Bonferroni correction on the alpha value.

## Results

### Functional network stability within and across behavioral states

We first assessed the similarity of infant functional networks during natural sleep and awake movie-watching (Figure 1A) by correlating the functional connectivity matrix of a single infant run with the average of all other infant runs. We found that infant functional networks were highly stable across participants within sleep (*M* = 0.681) and wake (*M* = 0.634) states, with slightly lower similarity across these two states (*M* = 0.610). Average similarity did not significantly differ between sleep and wake (difference *M* = 0.047, *p* = 0.074; Figure 1B). Within-state similarity was higher than across-state similarity for sleep (difference *M* = 0.070, *p* = 0.004), but did not reach significance for wake (difference *M* = 0.023, *p* = 0.280). These results were largely the same with a stricter motion threshold (0.2 mm, Figure S1A-D) and when runs were averaged within each unique participant prior to similarity calculation (Figure S2A-B). Thus, infant functional networks show comparable stability during sleep and wake states, despite the longer duration of rest runs compared to movie runs. Nonetheless, these networks have some differences that decrease similarity between states.

**Figure 1.**
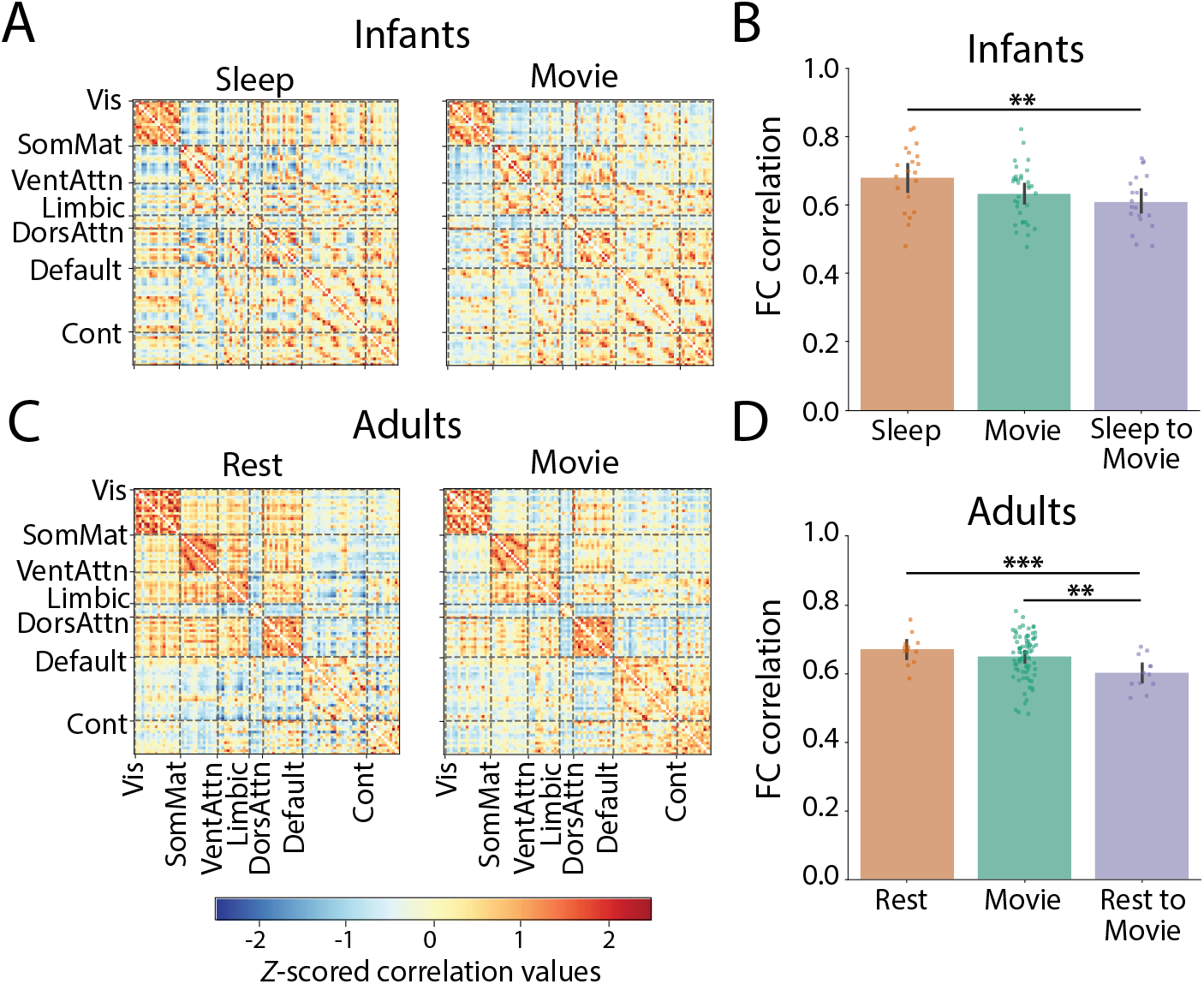
Functional connectivity in infants and adults in different behavioral states. (A,C) Visualization of the average functional connectivity matrix in infants and adults, respectively. (B, D) Similarity values comparing the upper triangle of the correlation matrix for an individual run with the average of all other runs within the infant and adult age groups, respectively. Dots represent individual run data and error bars represent 95% bootstrap confidence intervals. *** *p* <0.001, ** *p* <0.01. Network labels: visual (Vis), somatomator (SomMat), ventral attention (VentAttn), limbic, dorsal attention (DorsAttn), default, and frontoparietal control (Cont).

We next performed parallel analyses in adults as a point of comparison for the infant data (Figure 1C). Adult functional networks were highly stable across runs in rest (*M* = 0.671) and movie (*M* = 0.650) states, with slightly lower similarity across these two states (*M* = 0.602). Average similarity did not significantly differ between restand movie (difference *M* = 0.021, *p* = 0.168; Figure 1 D). Within-state similarity was higher than across-state similarity for both rest (difference *M* = 0.069, *p* <0.001) and movie (difference *M* = 0.048, *p* = 0.002). Importantly, the range and pattern of stability values for adults was almost identical to that of the infants, confirming the quality of the infant data and the reliability of functional networks early in development.

### Classification and generalization of behavioral state

Given the within versus across state differences observed above, we hypothesized that it should be possible to decode infant behavioral state from patterns of functional connectivity using multivariate classification (Lewis-Peacock and Norman, 2014). We first attempted to decode behavioral state within group based on whole-brain patterns of functional connectivity (Figure 2A). Indeed, we could robustly decode infant state (sleep vs. movie: *M* = 90.67% vs. 50%, *p* <0.001) and adult state (rest vs. movie: *M* = 95.56%, *p* <0.001). To unpack these whole-brain results, we next attempted to decode behavioral state in each group at the level of networks (Figure 2B). For both infants and adults, behavioral state was encoded throughout the brain, with functional connectivity varying by state within and across most pairs of networks. These results persisted if rather than subsampling participants to balance training examples we instead used a stratification approach that retained the original distribution of participants (Figure S3).

**Figure 2.**
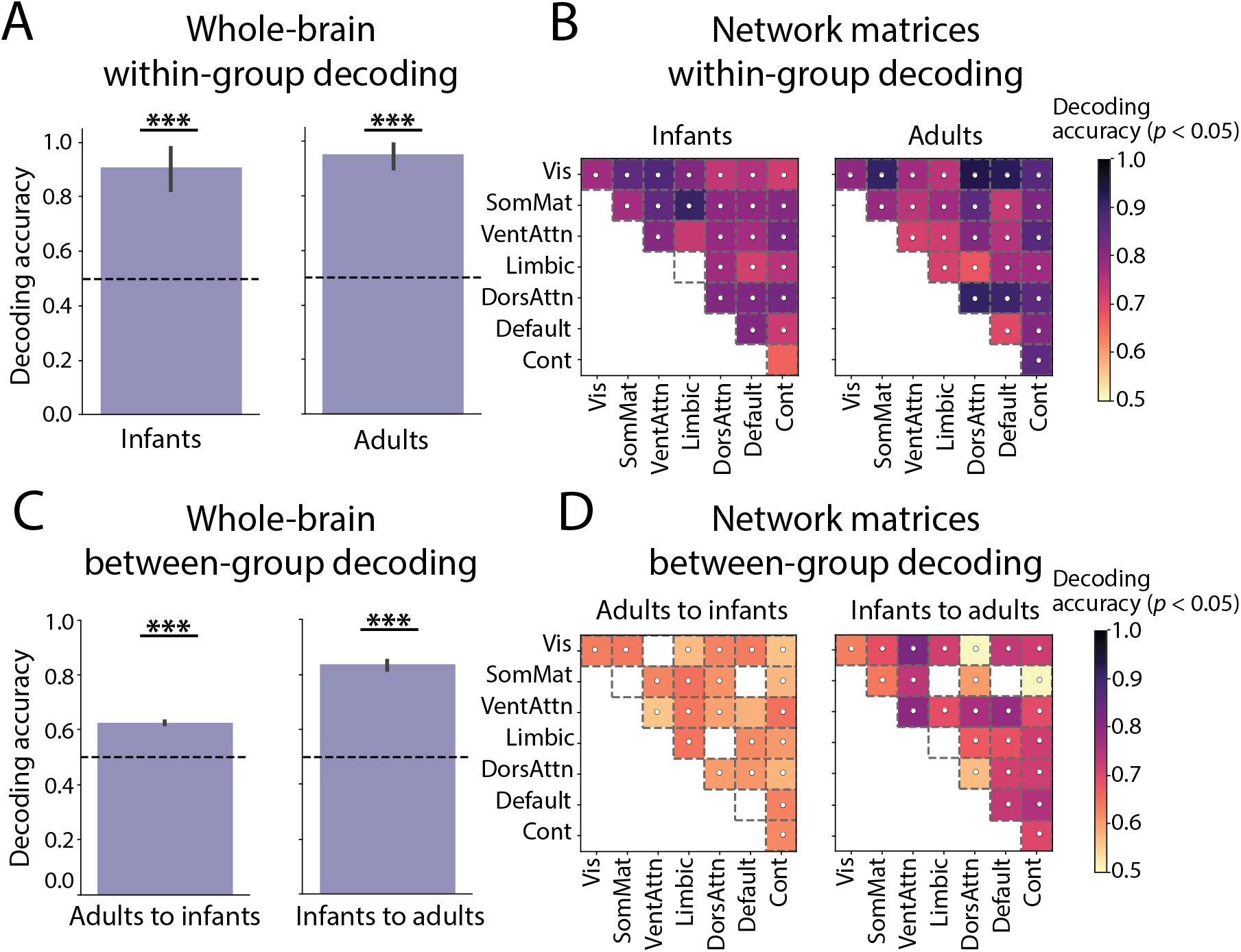
Classification and generalization of behavioral state based on functional connectivity. (A) Significant decoding of states across the whole brain within age groups compared with chance (0.5). (B) This was reflected in significant state decoding within and across pairs of networks (*p* <0.05, uncorrected), with darker cells indicating larger effects, and white dots denoting cells that survived Holms-Bonferroni correction. (C) Although infant state could be decoded with the adult classifier, adult state was decoded much more robustly with the infant classifier. (D) Repeating this analysis for networks revealed some generalization from adults to infants, but more robust and widespread generalization from infants to adults. Error bars represent variability across 10 folds. *** *p* <0.001

This evidence that behavioral state can be decoded within both infants and adults, and in similar networks, leaves open a question about whether the same connectivity features encode state information in each group. If so, the parcel-parcel connection weights learned by the state classifier in one group should enable decoding of state in the other group (Figure 2C). We first trained a classifier to distinguish rest versus movie in adults and tested whether it could generalize to distinguish sleep versus movie in infants (where rest and sleep were coded as the same class). This adult classifier was able to decode infant state reasonably well (*M* = 62.59% vs. 50%, *p* <0.001). Surprisingly, however, generalization worked even better in the opposite direction: A classifier trained to distinguish sleep versus movie in infants robustly decoded rest versus movie in adults (*M* = 83.56%, *p* <0.001). One interpretation of this asymmetry is that the connectivity features most useful for decoding infant state are represented in the adult brain, but that they are not the most useful features for adult decoding and thus are not weighted in the adult classifier in a way that would allow for their detection when tested on infant data. We again performed a network-level analysis to gain a better understanding of where this generalization occurs in the brain (Figure 2D). Although adult-to-infant generalization occurred in many network connections, infant-to-adult generalization was much stronger, particularly in visual-ventral attention connections and ventral attention within network connections.

### Functional network similarity between infants and adults

Infants and adults showed a comparable range of functional network similarity values within and across states in their own age group. Furthermore, some state-related features of functional connectivity were shared between infants and adults in the classifier generalization analyses. Here we test the similarity of infant and adult functional networks in more detail. In particular, we correlated the functional connectivity matrix of an infant in either the sleep or wake state with the average of all adult runs in either the rest or movie state.

Infants in both states had moderately similar functional connectivity to adults (Figure 3A), with no main effect of infant state (mean difference in infant sleep vs. infant movie: *M* = 0.023, CI = [−0.045, 0.049], *p* = 0.938). However, there was a main effect of adult state on similarity to infants (mean difference in adult rest vs. adult movie: *M* = −0.231, CI = [−0.270, −0.175], *p* <0.001). Follow-up tests revealed that, regardless of infant state, infants were more similar to adults watching a movie than adults resting (infant sleep to adult rest vs. infant sleep to adult movie: *M* = −0.099, *p* <0.001; infant movie to adult rest vs. infant movie to adult movie: *M* = −0.124, *p* <0.001; infant sleep to adult rest vs. infant movie to adult movie: *M* = −0.111, *p* <0.001; infant movie to adult rest vs. infant sleep to adult movie: *M* = −0.113, *p* <0.001). In other words, infant functional networks during sleep or wake better resemble adults performing a naturalistic viewing task than adults fixating a cross while resting. We did not find a significant interaction between infant state and adult state (*M* = −0.032, CI = [−0.072, 0.022], *p* = 0.334). Although infant sleep was numerically more similar to adult rest, and infant movie was numerically more similar to adult movie, there was no difference between infant sleep (*M* = 0.350) and infant movie (*M* = 0.337) in similarity to adult rest (difference *M* = 0.013, *p* = 0.490), nor a difference between infant sleep (*M* = 0.449) and infant movie (*M* = 0.461) in similarity to adult movie (difference *M* = −0.011, *p* = 0.498).

**Figure 3.**
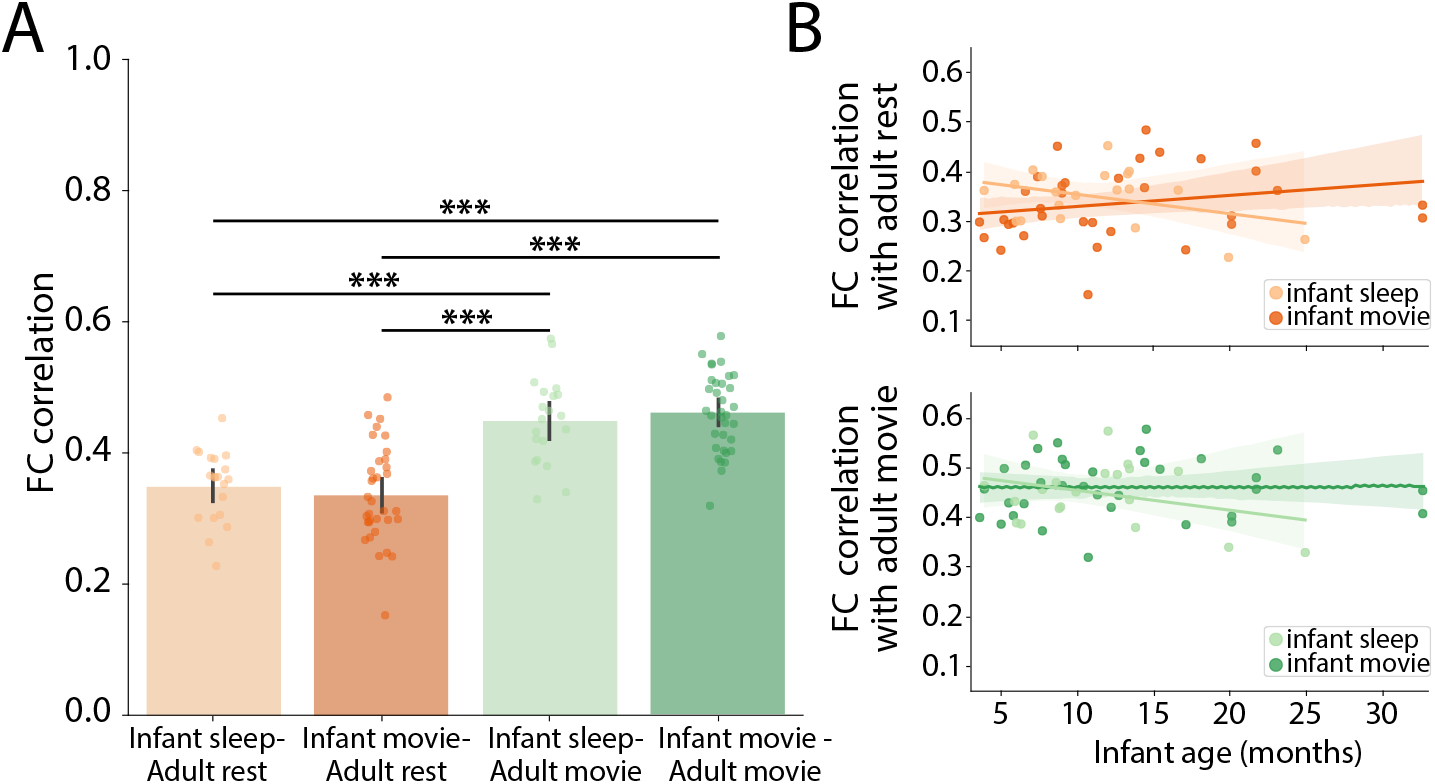
Functional network similarity between infants and adults. (A) Correlations between the upper triangle of individual infant functional connectivity matrices and the upper triangle of the average adult functional connectivity matrix, separated by different infant and adult states. Dots represent individual run data and error bars reflect 95% bootstrap confidence intervals. (B) Relationship between infant age and similarity to adults across the different comparisons. The shaded region signifies the 95% confidence interval for the line of best fit. *** *p* <0.001

The analyses above aggregated across all infants we tested. However, these infants ranged from 3 to 33 months old, spanning substantial developmental changes. We therefore performed an exploratory analysis of how similarity to adults changed with infant age. When compared to adult rest, the correlation between infant age and infant-adult similarity of functional connectivity did not reach significance for infant sleep (*r* = −0.366, *p* = 0.250) or infant movie (*r* = 0.228, *p* = 0.060). When compared to adult movie, there was again no significant correlation with age for infant sleep (*r* = −0.312, *p* = 0.340) or infant movie (*r* = 0.013, *p* = 0.926). Thus, there was no clear evidence of infant age-related differences in similarity to adults, though this question would be better addressed with a larger sample size and more uniform coverage of the age range.

### Network connections responsible for infant similarity to adult movie-watching

Our prior analysis showed a large main effect of adult state on similarity between infants and adults, with both sleeping and awake infants showing more similar functional connectivity to adults watching movies versus resting. To interpret this effect, we used a pointwise product approach to assess which parcel connections made a relatively larger contribution to this similarity. Averaging across infant sleep and awake states, the top connections responsible for similarity to adults were largely similar for adult rest and movie states. These connections were mainly between parcels labeled with the same network, particularly in the visual network and default-mode network (Figure 4A). To determine which connections contributed more to average infant similarity with adult movie compared with adult rest, we subtracted the average normalized pointwise product values of these two comparisons and calculated the significance of these differences at the network level with bootstrap resampling (Figure 4B). Whereas similarity between infants and adult rest was driven more by within-network connections, similarity between infants and adult movie was driven more by connections between the visual network and other networks. This suggests that movies reveal visual system interactions across the adult brain that are also present in the infant brain.

**Figure 4.**
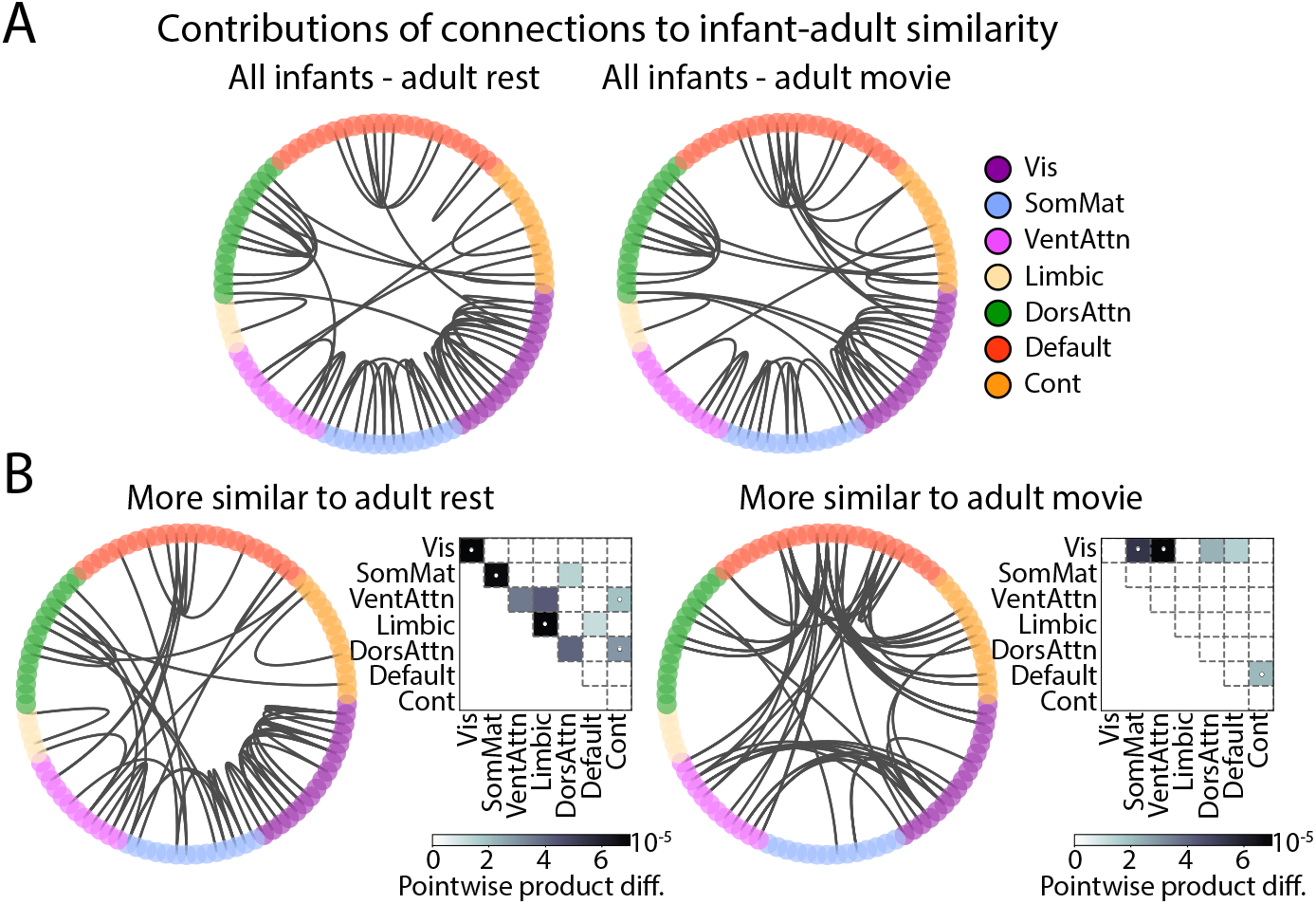
Contributions of individual network connections to infant-adult similarity, as assessed by normalized pointwise products. (A) Circos plots showing the top 1% of connections that contributed to the similarity between all infants and adult rest, and between all infants and adult movie. (B) Differences in the contribution of network connections to infant similarity with adult rest versus adult movie. Circos plots show the top 1% of differences. Matrices indicate which network connections contributed significantly more to infant similarity with adult rest and adult movie at *p* <0.05, uncorrected, with darker cells indicating larger effects. White dots indicate which network connections survived Holms-Bonferroni correction.

### Modulation of network connections driving adult similarity by infant state

The results so far show comparable overall levels of similarity between infant and adult functional connectivity for both sleep and wake infant states. However, which network connections are responsible for this overall adult similarity could differ by infant state. We again performed a normalized pointwise product analysis to test for differences in contributions to adult similarity between infant sleep and infant movie. Compared to adult rest (Figure 5A), similarity with infant sleep was driven by visual connections with other networks, control-somatomotor connections, and default-ventral attention connections, whereas similarity with infant movie was driven by ventral attention within-network connections, control within-network connections, control connections with other networks, and dorsal attention connections with other networks. This infant state-dependent reorganization of network structure similarity to adult rest persisted in the comparison to adult movie (Figure 5B). Similarity between infant sleep and adult movie was driven by visual-dorsal attention network connections, default connections to other networks, and control-somatomotor connections, whereas similarity between infant movie and adult movie was driven by control within-network connections and control connections to other networks. Thus, while infant state does not impact overall functional connectivity similarity to adults, it meaningfully changes which networks are more adult-like.

**Figure 5.**
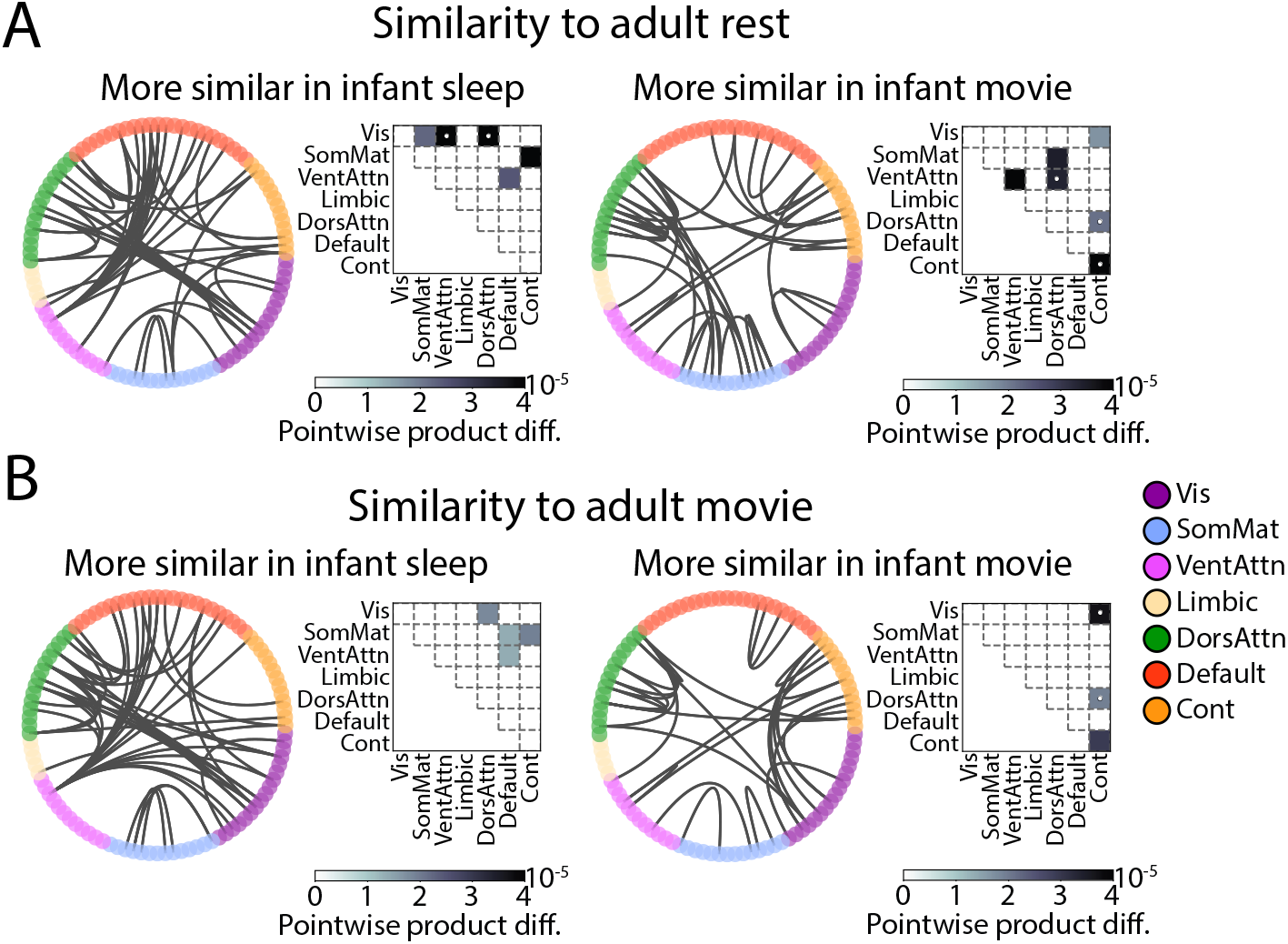
Contributions of individual network connections to infant-adult similarity dependent on infant state, as assessed by normalized pointwise product values. Circos plots show the top 1% of connections that contributed more to the labeled comparison. Matrices indicate which network connections contributed significantly more in that comparison (*p* <0.05, uncorrected), with darker cells indicating larger effects. White dots indicate which network connections survived Holms-Bonferroni correction. (A) Contributions of network connections to similarity with adult rest of infant sleep and infant movie. (B) Contributions of network connections to similarity with adult movie of infant sleep and infant movie.

## Discussion

In this study, we sought to fill a gap in the literature by comparing functional brain networks between sleeping and awake infants with fMRI. Whole-brain functional connectivity in infants differed between sleep and wake states, just as in adults. These states could be reliably decoded in held-out data within and across groups. A surprising asymmetry in across-group generalization suggests that infant state differences in functional connectivity are preserved in adults, but that the strongest indicators of adult state are not as relevant in infancy. Overall similarity with adult functional connectivity was comparable in both infant states (and stronger when adults watched movies vs. rested), but different networks contributed more or less to the overall level of similarity depending on state. This highlights the importance of considering infant behavioral state when assessing functional brain development relative to adults.

Differences in functional networks between rest and task are well-documented in adults and older children (Lynch et al., 2018; Sanchez-Alonso et al., 2021; Finn and Bandettini, 2021), but much less is known about how functional networks differ between infants who are asleep (as is most common in the literature; Zhang et al. 2019) versus awake and engaging in a task (Nielsen et al., 2023). Functional networks as measured with EEG differ between quiet and active sleep in neonates (Tokariev et al., 2019), and are more strongly coupled and less clustered during sleep than wake in 6-month-old infants (Smith et al., 2021). Our results expand on these findings in two ways: (1) by using fMRI, the dominant method to identify networks across the whole brain including away from the cortical surface; and (2) by using movies in the awake state, because it will be necessary for future awake infant fMRI studies to employ engaging stimuli to reduce head motion and ensure compliance for more than a few seconds.

We did not find support for our initial hypothesis of higher similarity between infants and adults in the awake state. Instead, infant functional networks were equally similar to adult functional networks overall, regardless of sleep/wake state. This result suggests that infant consciousness might not have a large impact on overall measures of functional network maturity, which bodes well for large-scale data collection efforts of infant functional connectivity that are relying entirely on sleeping fMRI data until early childhood (Fitzgibbon et al., 2020; Eyre et al., 2021; Volkow et al., 2021). Nevertheless, network-level analyses revealed modulation by infant state of which networks are most similar to adults, with connections between ventral attention to default mode networks among the strongest in infant sleep and connections within frontoparietal control network among the strongest in infant movie. These results are important because they mean that the functional maturity of certain networks may be underestimated in infant sleep data alone — particularly the frontoparietal control network, which is often characterized as one of the slowest developing networks in infancy (Gao et al., 2017; Zhang et al., 2019; Hu et al., 2022). Whether this is true in general or an artifact of sleep data will now require further investigation and data collection in awake infants.

Interestingly, infant functional networks were more similar to adults watching movies than to adults in a resting state, even when the infants were asleep. As shown in Figure 1, functional connectivity during adult rest consisted of stronger between-network connections than during adult movie, reflecting high integration (Bassett and Sporns, 2017). Although previous work has shown greater between-network connections during tasks compared to rest, this is not always the case for movies (Gonzalez-Castillo and Bandettini, 2018). Resembling adult movie, infants in both sleep and movie states showed weaker between-network connections, reflecting functional segregation. This observation may help explain the lower generalization from adults to infants in our multivariate decoding results: A classifier trained to distinguish adult rest and movie that then encounters infant data may choose the label “movie” more often if relying on features of network segregation. Thus, although some features relevant to state decoding may be preserved across development, those most heavily weighted in adults may not be applicable to infants. This result, which indicates that there are shared neural features between infants and adults despite a different neural pattern overall, fits with our prior work showing that infant neural event patterns during movie-watching are reflected in the adult brain, despite differences in optimal neural event timescales (Yates et al., 2022).

We used movies as a comparison to sleep for this study because they are rich, activate multiple networks across the brain, and are highly engaging to infants. Nonetheless, infant-adult network similarity may differ in awake rest or other tasks, such as those that tax attention or have an auditory component. Indeed, this is a key motivation of this research: the nature and kind of the functional networks identified in infants will likely depend on the state of the infant during data collection. In future work, it will be important to investigate the reorganization of functional networks within (Yin et al., 2020) and across tasks. We also did not find evidence of age-related changes in the similarity of functional networks between infants and adults in the current study. fMRI activity synchronizes across infants (and adults) watching the same movie (Yates et al., 2022), so this may not be the optimal task for characterizing individual differences. At the same time, infant functional connectivity during sleep only yields moderate identification rates within-session (Wang et al., 2021) and poor identification rates across sessions (Dufford et al. 2021; though see King et al. 2022 for discussion of methodological confounds in infant connectome fingerprinting). Thus, as the field of awake infant fMRI grows, it will be important to converge on the best task(s) for constructing reliable, yet individually identifiable functional networks.

As with all awake infant fMRI studies, there are a number of limitations to report. First, the amount of data used for calculating functional connectivity per participant was small (around 4 minutes), especially when compared to current recommendations (Birn et al., 2013; Noble et al., 2019). This was a practical necessity, given the difficulty of collecting data from infants, although likely influences power and reliability. Additionally, although we used stricter inclusion criteria than in previous studies with awake infants, our motion threshold was rather liberal. Importantly, infants and adults did not differ in the number of usable datapoints and our main results hold when we use a more standard motion threshold even on the awake data (0.2 mm; Figure S1). Nonetheless, the exclusion rate was much higher for infant movie runs (34/87 runs included) than for infant sleep runs (20/21 included), given that average motion is higher in awake than sleeping infants. In fact, given that motion hurts measurement of functional connectivity (Van Dijk et al., 2012; Power et al., 2012; Satterthwaite et al., 2012), our results may *underestimate* the impact of awake state on functional networks. Another limitation is that we treated adult rest and infant sleep as equivalent, despite known differences between rest and sleep in adults (Tagliazucchi and Laufs, 2014). Comparisons between infant movie and adult movie are better matched. In future work, it will be important to compare awake and sleeping infants to sleeping adults to fully understand sleep/wake differences across age groups. Characterizing sleep stages through simultaneous fMRI-EEG (Poppe et al., 2021) would also allow for a more fine-grained analysis of functional networks across different states. Finally, all of our analyses relied on an adult functional parcellation atlas (Schaefer et al., 2018), which may or may not be appropriate for delineating regions in the infant brain. Infant atlases are available (Scheinost et al., 2016; Oishi et al., 2019), although their use would complicate comparisons to adults for which they are not appropriate. Furthermore, it is unclear whether it is appropriate to import and label adult networks in the infant brain. For example, infant regions in the anatomical vicinity of the adult frontoparietal control network contribute more to adult similarity when infants are awake, but we do not yet know their function(s) (e.g., whether they are involved in executive control as in adults; Dosenbach et al. 2006). In fact, we previously showed that attention engages frontal cortex in infancy, but not as much parietal cortex (Ellis et al., 2021).

In conclusion, infants show distinctive functional network profiles during wake and sleep states that result in comparable overall similarity to adults driven by different networks. This highlights the added value of awake fMRI, in complement to more ubiquitous sleeping fMRI, for revealing early brain development.

## Acknowledgments

We are thankful to the families of infants who participated. We also acknowledge the hard work of the Yale Baby School team, including L. Rait,J. Daniels, A. Letrou, and K. Armstrong for recruitment, scheduling, and administration, and L. Skalaban, A. Bracher, D. Choi, andJ.Trach for help in infant fMRI data collection. Thank you to J. Wu, J. Fel, and A. Klein for help with gaze coding, R. Watts for technical support, and A. Holmes for analysis advice. We are grateful for internal funding from the Department of Psychology and Faculty of Arts and Sciences at Yale University. T.S.Y was supported by NSF Graduate Research Fellowship, Grant/Award Number: DGE 1752134. N.B.T-B. was further supported by the Canadian Institute for Advanced Research and the James S. McDonnell Foundation (https://doi.org/10.37717/2020-1208).

## Supplementary Data

**Table S1.** Demographic information for infant participants. ‘ID’ is a unique infant identifier (i.e., sXXXX_Y_Z), with the first four digits (XXXX) indicating the family, the fifth digit (Y) the child number within family, and the sixth digit (Z) the session number with that child. ‘Age’ is recorded in months. ‘Sex’ is female or male. ‘TR num’ is the total number of TRs recorded during that functional run. TR prop motion’ is the proportion of TRs included after motion exclusion. ‘TR prop eye’ is the proportion of TRs included after eye closure exclusion. ‘Eye IRR’ is the proportion of frames coded the same way across gaze coders. ‘Num coders’ is the number of gaze coders. ‘Movie’ is the movie that the infant watched.

| Infant sleep participants |  |  |  |  |  |
| --- | --- | --- | --- | --- | --- |
| ID | Age | Sex | Functional run | TR num | TR prop motion |
| s5927_1_1 | 3.90 | M | functional03b | 152 | 1.00 |
| s2067_1_2 | 5.90 | M | functional03b | 154 | 1.00 |
| s2986_1_1 | 6.00 | F | functional02c | 94 | 1.00 |
| s0307_1_1 | 6.30 | M | functional04b | 63 | 1.00 |
| s9986_1_1 | 7.10 | F | functional03b | 151 | 1.00 |
| s2057_1_2 | 7.70 | F | functional02c | 87 | 0.99 |
| s2986_1_2 | 8.60 | F | functional02b | 152 | 1.00 |
| s4107_1_1 | 8.80 | M | functional04 | 62 | 1.00 |
| s7286_2_2 | 8.90 | F | functional02b | 158 | 1.00 |
| s6057_1_4 | 9.90 | M | functional03 | 152 | 1.00 |
| s7286_2_3 | 11.80 | F | functional03b | 157 | 1.00 |
| s9986_1_4 | 12.00 | F | functional02b | 150 | 1.00 |
| s8687_2_2 | 12.60 | F | functional03 | 158 | 1.00 |
| s9986_1_5 | 13.30 | F | functional01b | 152 | 1.00 |
| s3097_1_4 | 13.40 | F | functional06 | 53 | 1.00 |
| s3097_1_4 | 13.40 | F | functional05 | 147 | 0.99 |
| s8187_1_4 | 13.80 | F | functional03 | 179 | 1.00 |
| s2097_1_4 | 16.60 | M | functional02b | 150 | 1.00 |
| s2307_1_1 | 19.90 | M | functional05b | 61 | 0.97 |
| s6057_1_7 | 24.90 | M | functional01b | 151 | 1.00 |
| Avg. | 11.24 | n/a | n/a | 129.15 | 0.99 |

**Table S1.** Demographic information for infant participants.
| Infant movie participants |  |  |  |  |  |  |  |  |  |
| --- | --- | --- | --- | --- | --- | --- | --- | --- | --- |
| ID | Age | Sex | Functional run | TR num | TR prop motion | TR prop eye | Eye IRR | Num coders | Movie |
| s6057_1_1 | 3.60 | M | functional03 | 90 | 1.00 | 1.00 | 1.00 | 2.00 | Aeronaut |
| s3097_1_1 | 3.90 | F | functional02a | 90 | 0.99 | 0.86 | 0.95 | 2.00 | Aeronaut |
| s6687_1_1 | 5.00 | F | functional02 | 71 | 1.00 | 0.71 | 0.89 | 3.00 | Mickey |
| s7017_1_1 | 5.20 | F | functional02b | 90 | 1.00 | 1.00 | 0.99 | 2.00 | Aeronaut |
| s4047_1_1 | 5.50 | F | functional02a | 90 | 1.00 | 1.00 | 0.94 | 2.00 | Aeronaut |
| s8687_1_2 | 5.80 | F | functional02 | 71 | 1.00 | 1.00 | 0.95 | 3.00 | Mickey |
| s2037_1_2 | 6.50 | M | functional03a | 90 | 0.99 | 0.98 | 0.92 | 2.00 | Aeronaut |
| s7017_1_2 | 6.60 | F | functional01b | 90 | 1.00 | 0.93 | 0.92 | 2.00 | Aeronaut |
| s6607_1_2 | 7.40 | M | functional02 | 90 | 1.00 | NaN | NaN | NaN | Aeronaut |
| s6057_1_2 | 7.60 | M | functional02a | 90 | 1.00 | 0.96 | 0.94 | 2.00 | Aeronaut |
| s2057_1_2 | 7.70 | F | functional02a | 90 | 0.97 | 0.81 | 0.92 | 2.00 | Aeronaut |
| s5037_1_1 | 8.70 | M | functional02b | 180 | 0.98 | 0.94 | 0.99 | 2.00 | Aeronaut |
| s0057_1_3 | 9.00 | F | functional02b | 90 | 1.00 | 0.96 | 0.95 | 2.00 | Aeronaut |
| s0687_1_2 | 9.00 | F | functional02 | 71 | 1.00 | 0.94 | 0.97 | 2.00 | Mickey |
| s7067_1_3 | 9.20 | F | functional01b | 90 | 1.00 | 0.98 | 0.98 | 2.00 | Aeronaut |
| s3607_1_2 | 10.40 | F | functional03 | 90 | 1.00 | 0.70 | 0.86 | 3.00 | Aeronaut |
| s8687_2_1 | 10.70 | F | functional02c | 90 | 1.00 | 0.96 | 0.98 | 2.00 | Aeronaut |
| s8687_1_3 | 11.00 | F | functional02d | 90 | 0.99 | 1.00 | 0.93 | 3.00 | Aeronaut |
| s6687_1_3 | 11.30 | F | functional01b | 90 | 1.00 | 0.97 | 0.90 | 2.00 | Aeronaut |
| s8037_1_2 | 12.20 | F | functional01c | 90 | 0.99 | 0.96 | 0.94 | 2.00 | Aeronaut |
| s0607_1_2 | 12.70 | M | functional01 | 90 | 1.00 | 0.98 | 0.92 | 2.00 | Aeronaut |
| s4607_1_5 | 14.10 | F | functional02 | 90 | 1.00 | 0.93 | 0.87 | 2.00 | Aeronaut |
| s1607_1_4 | 14.40 | M | functional03 | 90 | 0.98 | 0.93 | 0.91 | 3.00 | Aeronaut |
| s8687_1_4 | 14.50 | F | functional01a | 90 | 1.00 | 0.88 | 0.92 | 2.00 | Aeronaut |
| s6687_1_4 | 15.40 | F | functional01a | 90 | 1.00 | 0.84 | 0.86 | 2.00 | Aeronaut |
| s8687_1_5 | 17.10 | F | functional02a | 90 | 1.00 | 0.99 | 0.98 | 2.00 | Aeronaut |
| s6687_1_5 | 18.10 | F | functional01c | 90 | 1.00 | 0.97 | 0.92 | 2.00 | Aeronaut |
| s8687_1_7 | 20.10 | F | functional02 | 90 | 1.00 | 1.00 | 0.99 | 2.00 | Aeronaut |
| s6687_1_6 | 20.10 | F | functional02b | 90 | 1.00 | 0.99 | 0.99 | 2.00 | Aeronaut |
| s2307_1_2 | 21.70 | M | functional02 | 71 | 1.00 | 1.00 | 0.98 | 6.00 | Mickey |
| s2307_1_2 | 21.70 | M | functional04 | 71 | 1.00 | 1.00 | 0.97 | 6.00 | Mickey |
| s8187_1_8 | 23.10 | F | functional01 | 142 | 0.99 | 0.99 | 0.98 | 2.00 | Mickey |
| s0307_2_1 | 32.60 | M | functional01 | 71 | 1.00 | 0.63 | 0.93 | 2.00 | Mickey |
| s0307_2_1 | 32.60 | M | functional03 | 71 | 1.00 | 0.82 | 0.77 | 2.00 | Mickey |
| Avg. | 12.78 | n/a | n/a | 90.26 | 0.99 | 0.93 | 0.94 | 2.39 | n/a |

**Figure S1.**
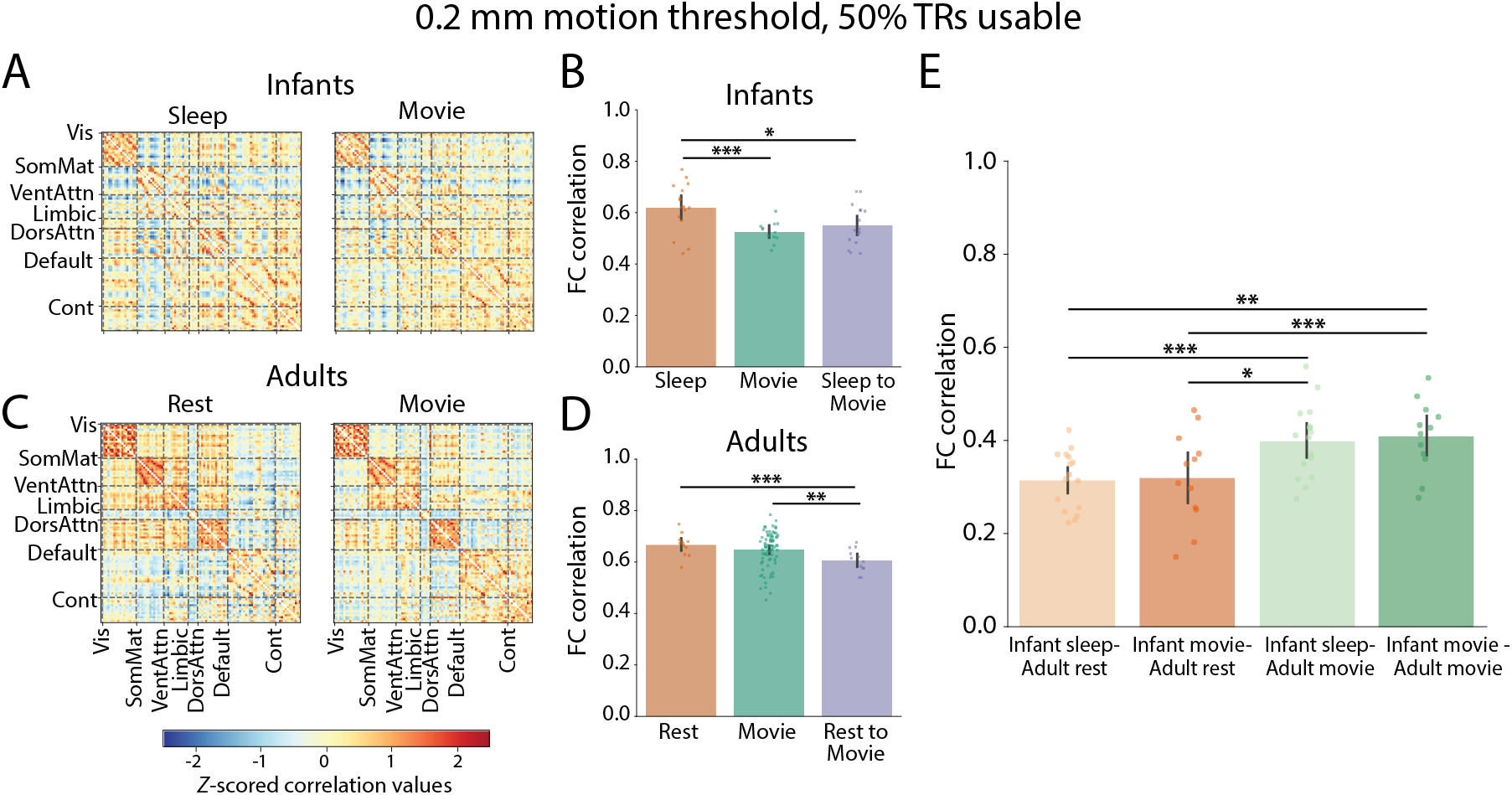
Functional connectivity in infants and adults in different states, using a stricter motion threshold (0.2mm, with 50% of TRs needing to be usable). (A,C) Visualization of the average functional connectivity matrix in infants and adults, respectively. Infant data was reduced to 16 infant sleep runs (out of 20) and 12 infant movie runs (out of 34). Adult movie runs were reduced to 76 (out of 78), while all adult rest runs were retained. (B, D) Correlation values comparing the upper triangle of the correlation matrix for an individual run with the average of all other runs. (E) Correlations between the upper triangle of individual infant functional connectivity matrices and the upper triangle of the average adult functional connectivity matrix, separated by different infant and adult states. Dots represent individual run data, and error bars reflect 95% bootstrap confidence intervals. *** *p* <0.001, ** *p* <0.01 * *p* <0.05. Network labels: visual (Vis), somatomator (SomMat), ventral attention (VentAttn), limbic, dorsal attention (DorsAttn), default, and frontoparietal control (Cont).

**Figure S2.**
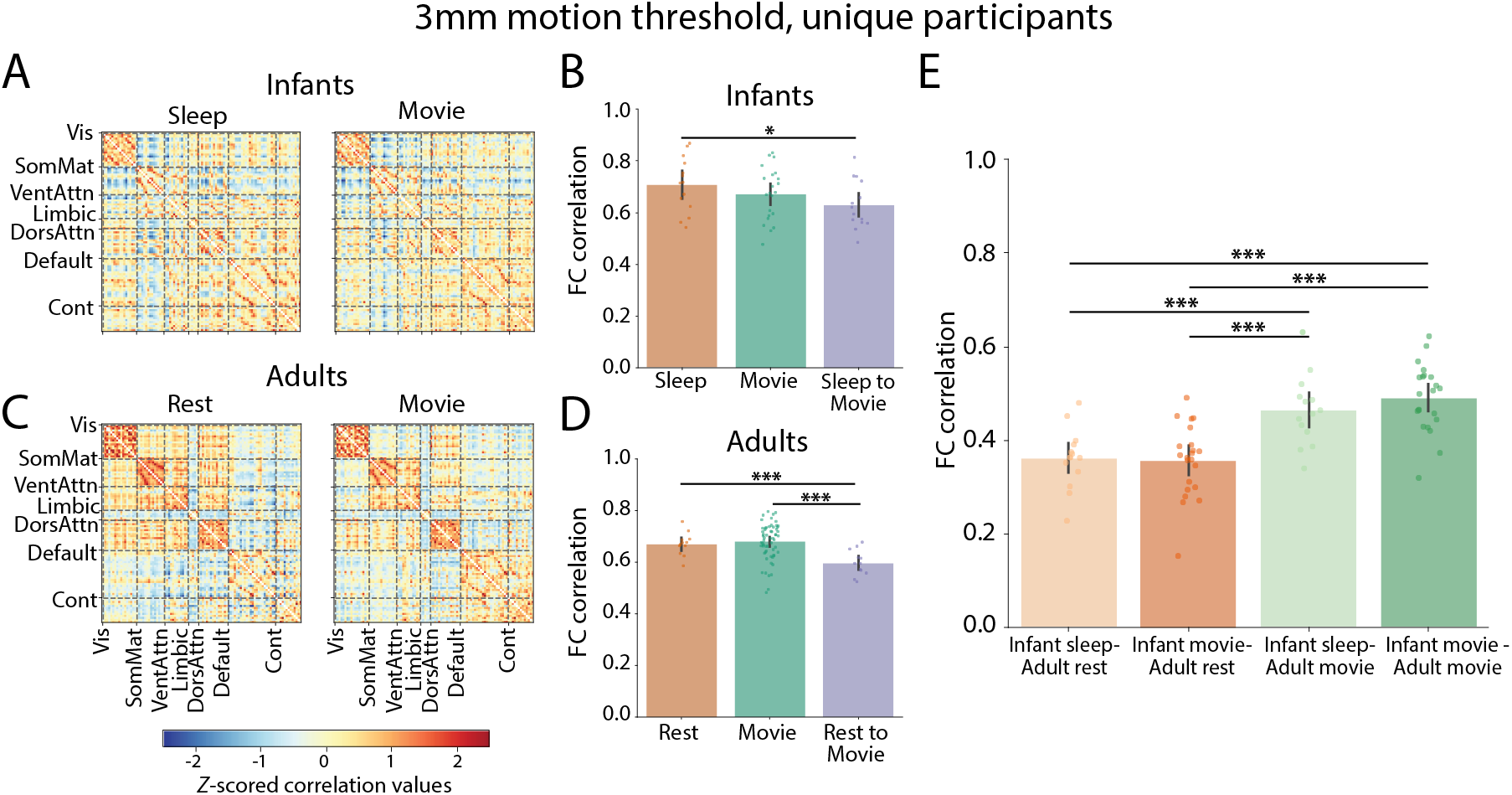
Functional connectivity in infants and adults in different states after averaging across runs within unique participants. (A,C) Visualization of the average functional connectivity matrix in infants and adults, respectively. Infant data reflect 14 unique sleep participants and 22 unique movie participants. Adult data reflect 12 unique rest participants and 60 unique movie participants. (B, D) Correlation values comparing the upper triangle of the correlation matrix for an individual participant with the average of all other participants. (E) Correlations between the upper triangle of individual infant functional connectivity matrices and the upper triangle of the average adult functional connectivity matrix, separated by different infant and adult states. Dots represent individual run data, and error bars reflect 95% bootstrap confidence intervals. *** *p* <0.001, * *p* <0.05.

**Figure S3.**
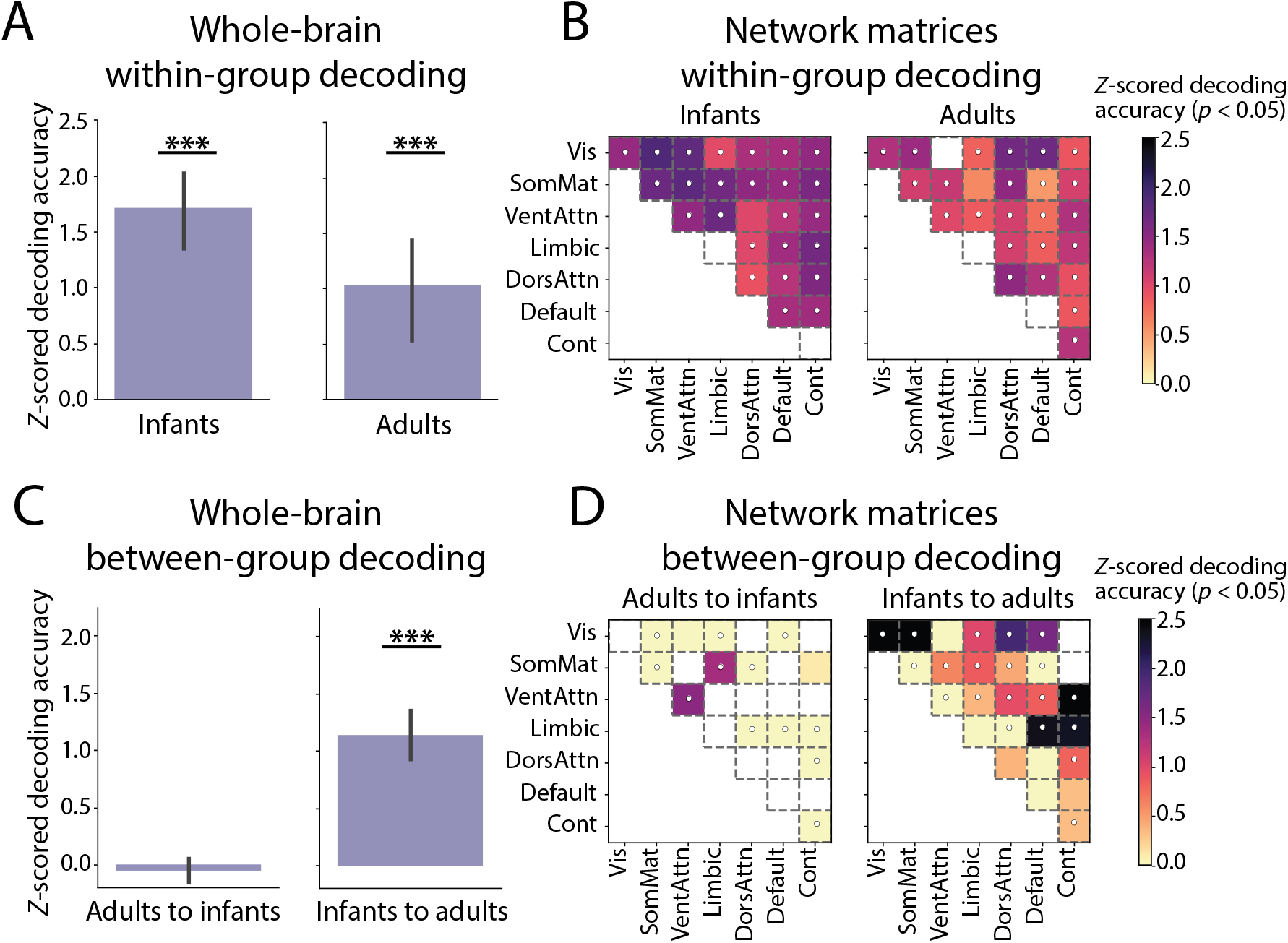
Classification and generalization of behavioral state based on functional connectivity, using a stratification approach to match the number of runs of each state in model training with the original proportion of participants in each state from the full dataset (in contrast to the subsampling approach used in the main text to equate the number of training examples between states). (A) Significant decoding of states across the whole brain within age groups compared with permuted null distributions generated by shuffling labels. (B) This was reflected in significant state decoding within and across pairs of networks (*p* <0.05, uncorrected, with white dots denoting significance after Holms-Bonferroni correction). (C) Infant state could not be decoded with the adult classifier, but adult state could be decoded with the infant classifier. This is mostly consistent with the main results showing lower generalization from adults to infants (though chance performance here vs. slightly above-chance performance with the subsampling approach). (D) Repeating this analysis at the level of networks revealed some generalization from adults to infants, but more robust and widespread generalization from infants to adults. Error bars represent variability across 10 folds. *** *p* <0.001.

**Figure S4.**
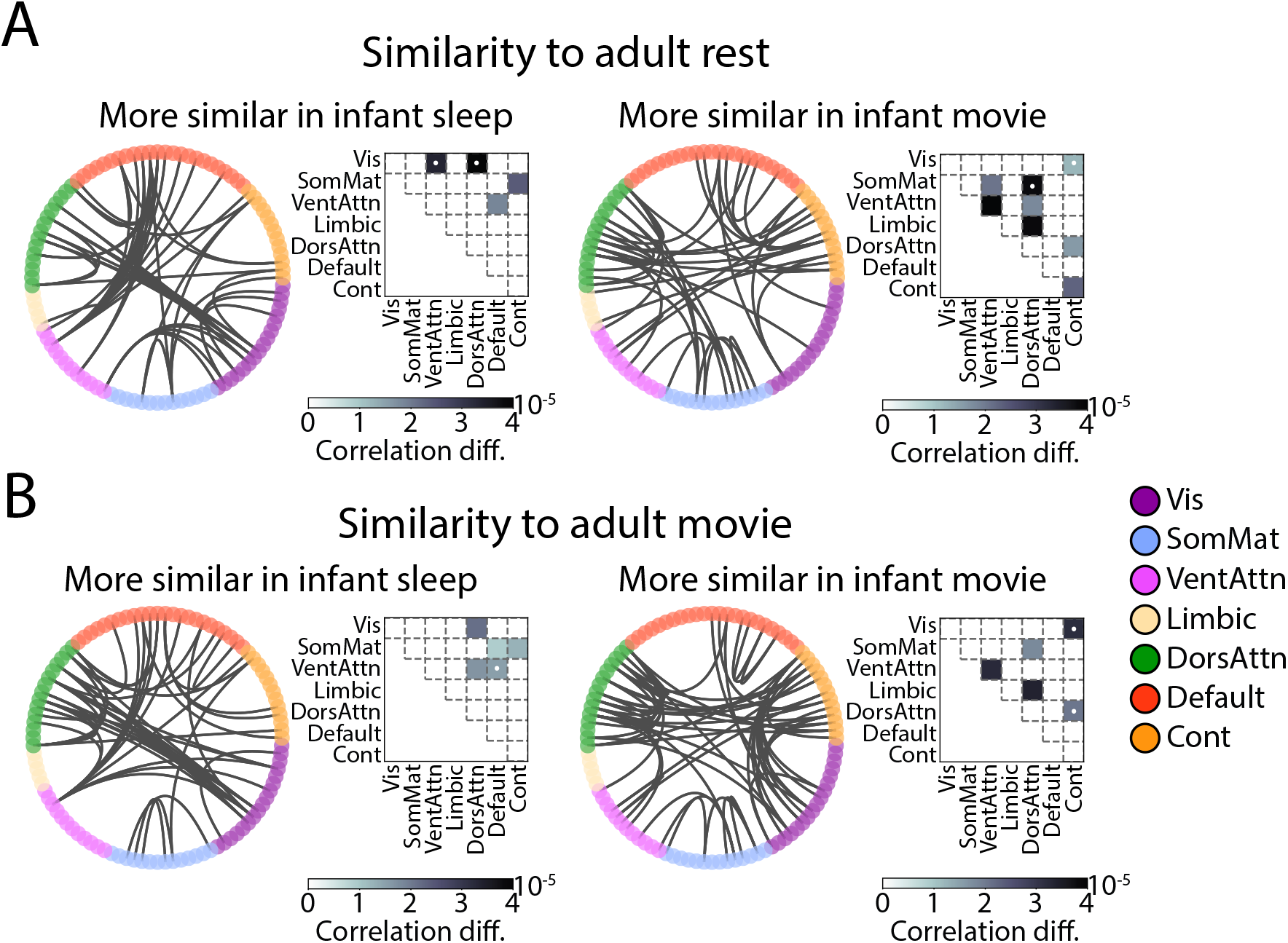
Contributions of individual network connections to infant-adult similarity, as assessed by lesioning connections and assessing the resulting change in correlation. Differences in the contribution of network connections to (A) similarity between infant sleep vs. infant movie to adult rest, and (B) similarity between infant sleep vs. infant movie to adult movie. Circos plots show the top 1% of differences. Matrices indicate which network connections contributed significantly more to infant sleep vs. movie similarity with adults at *p* <0.05, uncorrected, with darker cells indicating larger effects. White dots indicate which network connections survived Holms-Bonferroni correction.

